# Correspondence between gene expression and neurotransmitter receptor and transporter density in the human brain

**DOI:** 10.1101/2021.11.30.469876

**Authors:** Justine Y. Hansen, Ross D. Markello, Lauri Tuominen, Martin Nørgaard, Elena Kuzmin, Nicola Palomero-Gallagher, Alain Dagher, Bratislav Misic

## Abstract

Neurotransmitter receptors modulate the signaling between neurons. Thus, neurotransmitter receptors and transporters play a key role in shaping brain function. Due to the lack of comprehensive neurotransmitter receptor/transporter density datasets, microarray gene expression is often used as a proxy for receptor densities. In the present report, we comprehensively test the expression-density association for a total of 27 neurotransmitter receptors, receptor binding-sites, and transporters across 9 different neurotransmitter systems, using both PET and autoradiography imaging modalities. We find poor spatial correspondences between gene expression and density for all neurotransmitter receptors and transporters except four single-protein metabotropic receptors (5-HT_1A_, D_2_, CB_1_, and MOR). These expression-density associations are related to population variance and change across different classes of laminar differentiation. Altogether, we recommend using direct measures of receptor and transporter density when relating neurotransmitter systems to brain structure and function.

## INTRODUCTION

Neurotransmitter receptors and transporters support synaptic communication, regulating signal transmission from neuron to neuron. As such, regional variation of receptor and transporter distributions shapes the functional specialization of the brain [27, 34, 60, 79, 84, 91]. Recent studies have used receptor densities to tune computational models and have related specific receptors, as well as excitatory-inhibitory ratios, to neurodevelopment, cognition, neural dynamics, and disease [20, 39, 44, 64, 80]. However, due to the lack of comprehensive neurotransmitter receptor and transporter density datasets (open-source or otherwise), receptor/transporter densities are often substituted with microarray gene expression from the Allen Human Brain Atlas [37], under the assumption that levels of gene expression are correlated with cell surface protein abundance [6, 14–16, 20, 30, 33, 39, 44, 64, 65, 80].

Despite the frequent replacement of receptor/transporter densities with gene expression, the assumed correlation between gene expression and receptor/transporter density has yet to be comprehensively and formally tested across multiple neurotransmitter systems. Indeed, there are several reasons gene expression may not be correlated with receptor density. First, microarray gene expression measures the outcome of gene transcription and the abundance of mRNA, not levels of protein. Importantly, levels of mRNA and protein are often not correlated within the same tissue [52, 76]. Second, several steps are involved between protein translation and expression of the receptor/transporter on the cell surface, including post-translational modifications, protein folding, and reaching a designated cellular target. Variations in the activity of these processes will affect receptor/transporter density. Third, multiple genes in the Allen Human Brain Atlas show high intersubject variability, indicating the possible unreliability of such group-averaged expression levels. Altogether, a comprehensive study mapping receptor/transporter densities and gene expression levels is necessary to determine whether neurotransmitter receptors and transporters show density-expression associations in the human cortex.

Here we investigate whether microarray gene expression can be used to estimate neurotransmitter receptor/transporter densities in the cortex. To measure gene expression levels, we use the Allen Human Brain Atlas that code for specific neurotransmitter receptors or transporters [37, 47]. Additionally, we use both positron emission tomography (PET)- and autoradiography-derived measures of neurotransmitter receptor densities for a total of 27 neurotransmitter receptors and transporters across 9 different neurotransmitter systems [1, 8, 10, 22–24, 31, 38, 41, 42, 56, 58, 59, 62, 71, 74, 82, 91]. To ensure results are not biased by methodological choices, we repeat our analyses in a separate parcellation resolution and use a conservative spatial autocorrelation-preserving null model [2, 49]. We find that measures of gene expression can generally not be used as a proxy for receptor nor transporter density except for a select few neurotransmitter receptors: 5-HT_1A_ (serotonin), D_2_ (dopamine), CB_1_ (cannabinoid), and MOR (opioid). Finally, we provide evidence that the link between receptor/transporter density and gene expression is related to inter-subject genetic variability.

## METHODS

All code and data used to perform the analyses can be found at https://github.com/netneurolab/hansen_gene-receptor.

### PET data acquisition

Volumetric PET images were collected for 18 different neurotransmitter receptors and transporters across 9 different neurotransmitter systems [1, 8, 10, 22–24, 31, 38, 41, 42, 56, 58, 59, 62, 71, 74, 82, 91]. To protect patient confidentiality, individual participant maps were averaged within studies before being shared. Each study, the associated receptor/transporter, tracer, number of healthy participants, age, and reference with full methodological details can be found in Table S1. In all cases, only scans from healthy participants were included. Images were acquired using best practice imaging protocols recommended for each radioligand. Altogether, the images are an estimate of receptor densities and we therefore refer to the measured value (i.e. binding potential, tracer distribution volume) simply as density PET images were all registered to the MNI-ICBM 152 nonlinear 2009 (version c, asymmetric) template, then parcellated to a parcellation with 68 and 219 cortical regions, as well as 15 subcortical regions, according to the Lausanne atlas [17, 21]. Receptors and transporters with more than one mean image of the same tracer (i.e. 5-HT_1B_, D_2_, mGluR_5_, and VAChT) were combined using a weighted average. Finally, each tracer map corresponding to each receptor/transporter was z-scored across regions and concatenated into a final region × receptor matrix of relative densities. This data was presented and used originally in [35].

### Autoradiography data acquisition

In vitro receptor autoradiography data were originally collected and processed as described in [91]. Fifteen neurotransmitter receptor densities across 44 cytoarchitectonically identified areas in three post-mortem brains were acquired from Supplementary Table 2 of [91] (see Table S2 for a complete list of receptors included in the autoradiography dataset). Note that GABAA and GABAA/BZ refer to the same receptor, but that GABAA/BZ refers specifically to GABAA receptors containing the allosteric benzodiazepine binding site, as opposed to receptors containing only the GABA neurotransmitter binding site. To best compare PET data analyses with the autoradiography dataset, a region-to-region mapping was manually created between the 44 available cortical regions in the autoradiography dataset and the 34 left hemisphere cortical Desikan Killiany regions. In only one case (the insula) was there no suitable mapping between the autoradiography data and the Desikan Killiany atlas. As such, the 44-region autoradiography atlas was converted to 33 Desikan Killiany left hemisphere regions. Finally, receptor densities were z-scored and averaged across laminar layers, to create a single map of receptor densities across the cortex.

### Microarray gene expression

Regional microarray expression data were obtained from six post-mortem brains provided by the Allen Human Brain Atlas (http://human.brain-map.org/) [37]. Since only two of the six brains included samples from the right hemisphere, main analyses were conducted on the left hemisphere only. All processing was performed using the *abagen* toolbox (https://github.com/netneurolab/abagen [47]). These data were processed and mapped to parcellated brain regions at 34 and 111 left hemisphere cortical grey matter nodes according to the Lausanne anatomical atlas [17, 21]. For completeness, data were also parcellated to 15 bilateral subcortical regions. Due to the coarse subcortical parcellation, sufficient probes were available for both hemispheres. We therefore show expression-density associations across 15 bilateral subcortical regions in Fig. S1 but due to the low number of observations (brain regions), we do not report nor interpret corrrelation coefficients or *p*-values.

Microarray probes were reannotated using data provided by Arnatkeviĉiūtė et al. [3]. A single microarray probe with the highest differential stability, Δ_*S*_ (*p*), was selected to represent each gene [36], where differential stability was calculated as:

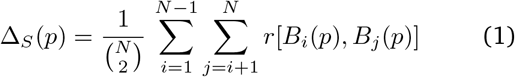

Here, *r* is Spearman’s rank correlation of the expression of a single probe p across regions in two donor brains, *B_i_* and *B_j_*, and *N* is the total number of donor brains. Differential stability is the average correlation across every pair of donor brains of a probe’s expression. This procedure retained 20 232 probes, each representing a unique gene.

Next, samples were assigned to brain regions using MNI coordinates generated via non-linear registrations (https://github.com/chrisfilo/alleninf) by finding the nearest region, up to 2 mm away. To reduce the potential for misassignment, sample-to-region matching was constrained by hemisphere and cortical/subcortical divisions [3]. If a brain region was not assigned any sample based on the above procedure, the sample closest to the centroid of that region was selected in order to ensure that all brain regions were assigned a value.

Inter-subject variation was addressed by normalizing tissue sample expression values for each donor across genes using a scaled robust sigmoid function [29]:

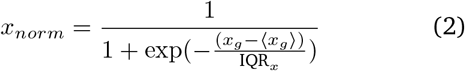

where 〈*x_g_*〉 is the median and IQR is the normalized interquartile range of the expression value of a single gene across regions. Normalized gene expression values were then rescaled to a unit interval:

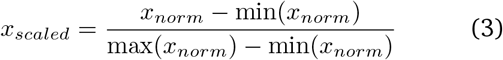

Gene expression values were normalized across tissue samples using the same procedure. Samples assigned to the same brain region were then averaged separately for each donor. Scaled regional expression profiles were finally averaged across donors.

### Gene-receptor pairs

With the notable exception of the GABA_B_ receptor, metabotropic neurotransmitter receptors are monomeric structures, and thus a single gene codes for the entire receptor. Therefore, the expression of the receptor-coding gene was correlated with the density of the receptor itself. The GABA_B_ and ionotropic receptors are characterized by being multimeric protein complexes, so each receptor was correlated with microarray expression of all possible receptor subunits. Below, we outline each multimeric case.

- GABA_A_ is a pentamer typically composed of three primary subunits (*α*_1_, *β*_2_, and *γ*_2_) but can be built out of a total of nineteen different subunits. For simplicity, we show results for the three primary subunits in the main text, but results for the remaining sixteen subunits can be found in Fig. S2.
- GABA_B_, a multimeric metabotropic receptor, is composed of two subunits. We show both in the main analyses.
- AMPA is a heterotetramer that typically consists of two pairs of duplicate subunits. These two pairs can be formed from any combination of four subunits. We show results for the gene whose expression is most highly correlated with AMPA density in the main text (*GRIA1*).
- NMDA is also a heterotetramer, typically composed of two *N1* and two *N2* subunits, although there are four different *N2*-encoding genes, as well as a third subunit (*N3*) for which there are two subunit-encoding genes. The main analyses use the expression of the *N1*-encoding gene (*GRIN1*).
- Kainate exists as both a homotetramer and heterotetramer, built from any of five subunits. We show results for the gene whose expression is most highly correlated with kainate density in the main text (*GRIK2*).
- *α*_4_*β*_2_ is a pentamer typically composed of two *α*_4_ subunits and three *β*_2_ subunits [19]. However, the ligand used for the autoradiograph, epibatidin, binds to any heteromeric nicotinic receptors that contain both an alpha subunit (*α*_2_–*α*_7_, *α*_9_, *α*_10_) and a beta subunit (*β*_2_–*β*_4_). The most abundant such receptor in the brain is the *α*_4_*β*_2_ receptor, so we focus on gene expression of the *α*_4_ and *β*_2_ subunits (*CHRNA4* and *CHRNB2*, respectively) in the main text.

Correlations between neurotransmitter receptor density and multiple subunit expression were corrected for multiple comparisons using the Benjamini-Hochberg FDR correction [11]. Correlation coefficients and corrected *p*-values (see *Null model*) for all subunits can be found in Supplementary Table 3 (PET) and Supplementary Table 4 (autoradiography).

### Null model

Spatial autocorrelation-preserving permutation tests were used to assess statistical significance of associations across brain regions, termed “spin tests” [2, 48]. Parametric *p*-values were not used because spatially embedded systems such as the brain violate the assumption that observations (brain regions) are independent from one another. We created a surface-based representation of the parcellation on the FreeSurfer fsaverage surface, via files from the Connectome Mapper toolkit (https://github.com/LTS5/cmp). We used the spherical projection of the fsaverage surface to define spatial coordinates for each parcel by selecting the coordinates of the vertex closest to the center of the mass of each parcel [85]. These parcel coordinates were then randomly rotated, and original parcels were reassigned the value of the closest rotated parcel (10 000 repetitions). Parcels for which the medial wall was closest were assigned the value of the next most proximal parcel instead. The procedure was performed at the parcel resolution rather than the vertex resolution to avoid upsampling the data, and only to the left hemisphere. In the autoradiography dataset, null correlations were computed ignoring the insula and regions resampled to the insula, for a maximum of three ignored brain regions.

## RESULTS

Using the group-averaged healthy control PET-derived neurotransmitter receptor and transporter densities, we find that in almost all cases, there is no significant correlation between receptor density and receptor gene expression (Fig. 1; for results in the subcortex see Fig. S1). Indeed, only five single-protein inhibitory metabotropic receptors (5-HT_1A_, D_2_, CB_1_, M_1_ and MOR) show significant relationships (*p*_spin_ < 0.05) with the expression of their corresponding genes (*HTR1A, DRD2, CRN1, CHRM1*, and *OPRM1*, respectively). Notably, consistent with previous reports, the GABAA receptor is positively correlated (*r* > 0.6) with the expression of its *β*_2_ subunit, and negatively correlated (*r* < –0.6) with the expression of its *α*_3_, *α*_5_, *β*_1_, and *γ*_1_ [53, 58], although these relationships are not significant after correcting for multiple comparisons (see Fig. S2 for scatter plots with remaining GABA_A_ subunits and Supplementary Table 3 for reported Pearson’s *r* and corrected *p*-values; [11]). Similarly, using the autoradiography dataset, we find that in almost all cases, gene expression is a poor approximator of neurotransmitter receptor density (Fig. 2). We again find a close correspondence between 5-HT1A and the expression of *HTR1A* (*r* = 0.82, *p*_spin_ = 0.002). However, unlike what was observed with the PET dataset, M_1_ does not show significant correlations with *CHRM1*. We additionally find a significant correlation between *α*_2_ receptor density and *ADRA2A* gene expression, although the effect is less apparent. No other neurotransmitter receptor besides 5-HT1A in the autoradiography dataset has a close correspondence between receptor density and gene expression.

**Figure 1.**
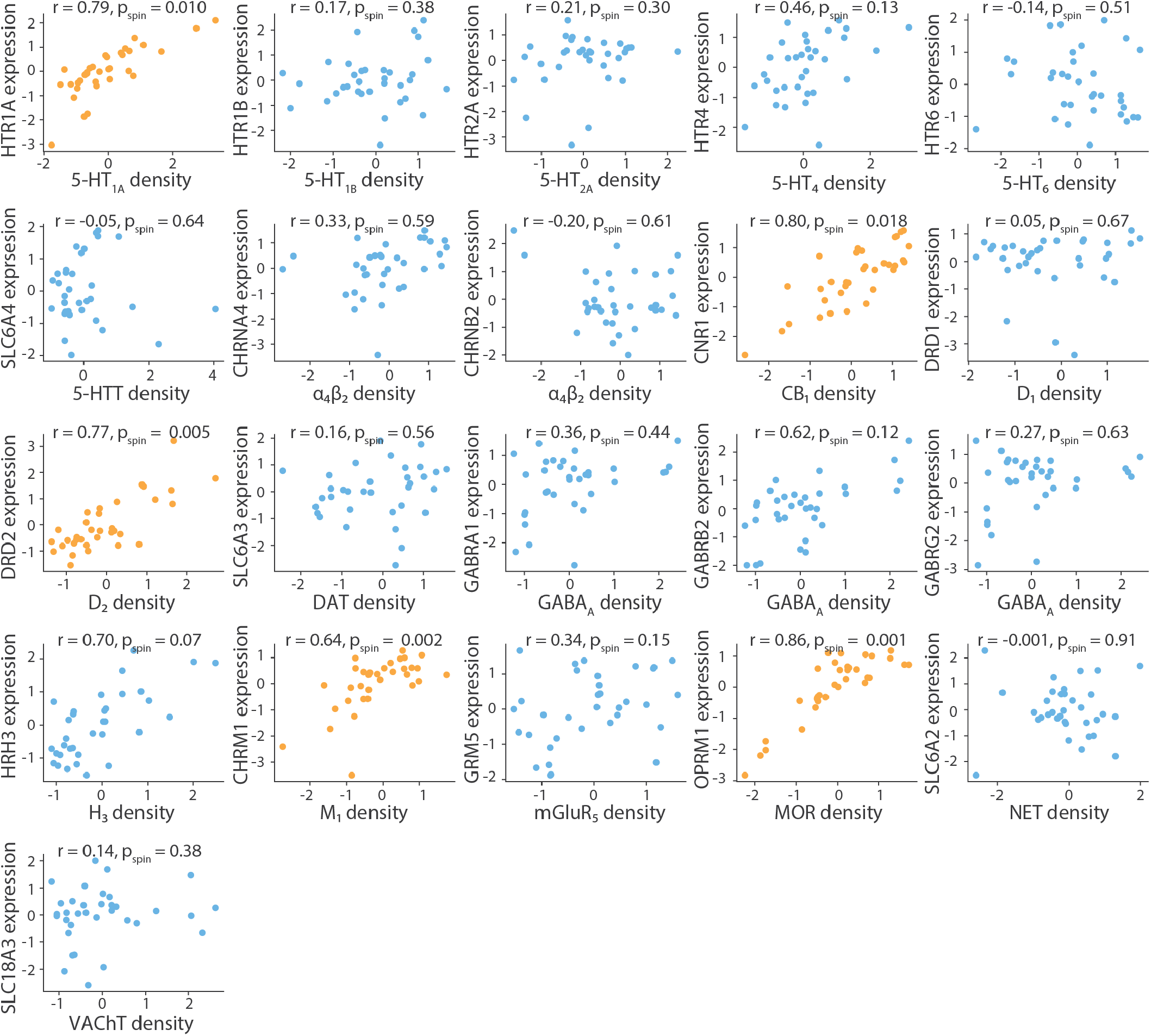
PET-derived receptor/transporter densities versus gene expression. PET tracer maps for 18 different neurotransmitter receptors and transports reveal that the density of only five neurotransmitter receptors correlates significantly with the expression of their corresponding gene, across 34 Desikan Killiany regions in the left cortex. Yellow scatter plots indicate significant density-expression correspondence, against an autocorrelation-preserving null model. All receptor densities are z-scored.

**Figure 2.**
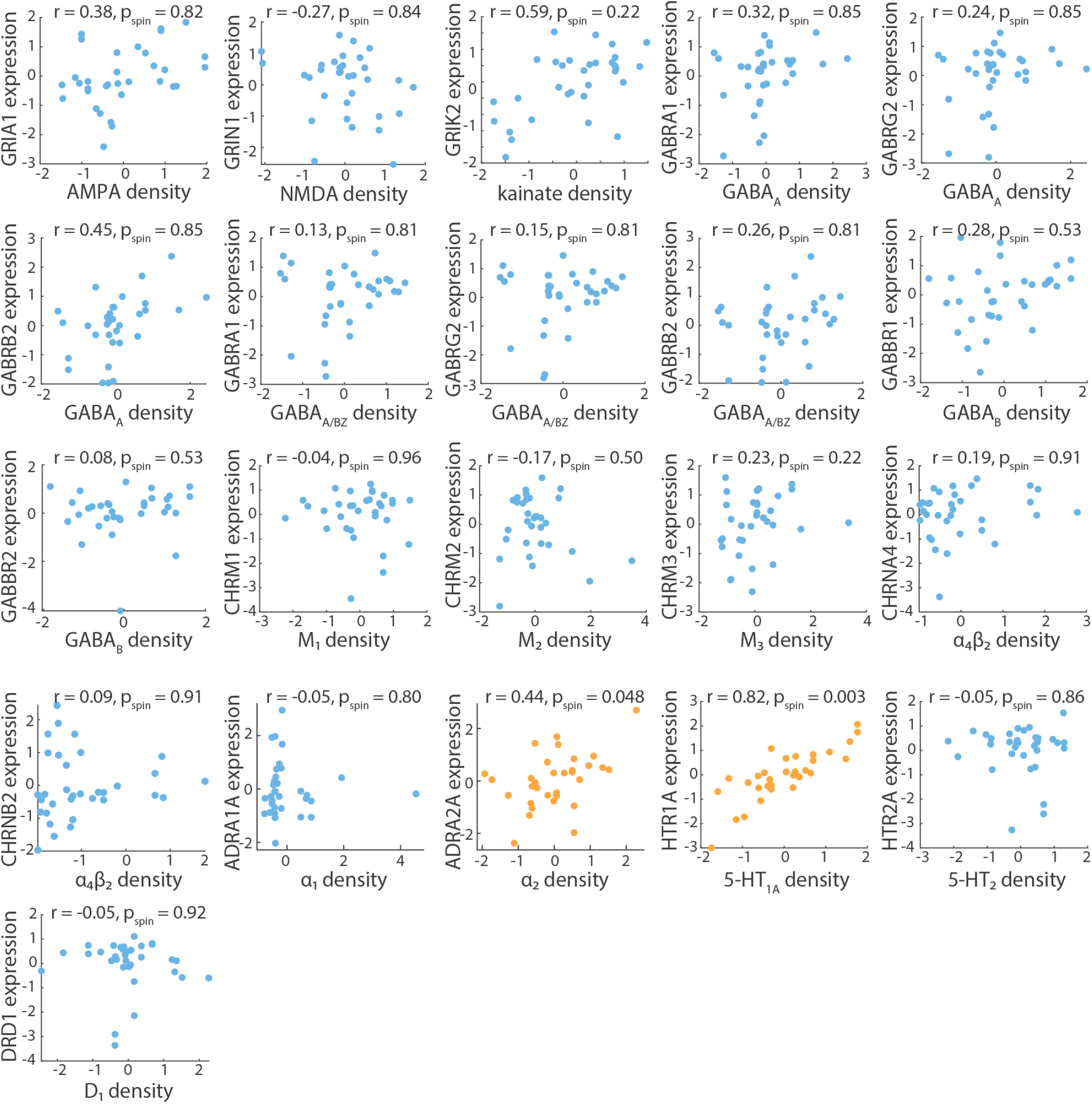
Autoradiography-derived receptor densities versus gene expression. Autoradiographs for 15 different neurotransmitter receptors reveal that only 5-HT_1A_ and *α*_2_ correlate significantly with the expression of their corresponding genes (*HTR1A* and *ADRA2A*, respectively), across 33 Desikan Killiany regions in the left cortex. Yellow scatter plots indicate significant densityexpression correspondence, against an autocorrelation-preserving null model. All receptor densities are z-scored.

To ensure results are not influenced by the choice of brain parcellation, we repeated the PET analyses in a finer parcellation of 111 left hemisphere cortical regions (Fig. S3; [17]). At this higher resolution, we find that 5-HT_1A_, D_2_, CB_1_, and MOR still show close correspondences with their associated genes, although CB_1_’s correlation has dropped to *r* = 0.66. M_1_ is still significantly correlated to *CHRM1* although to a lesser degree (*r* = 0.35, *p*_spin_ = 0.017). Furthermore, we find an additional significantly-correlated gene-receptor pair (*GRM5*–mGluR_5_: *r* = 0.44, *p*_spin_ = 0.008). Altogether, 5-HT_1A_, D_2_, CB_1_, and MOR show stable and high expression-density correspondence across both spatial scales and imaging modalities.

We next sought to understand why certain neurotransmitter receptors demonstrate expression-density correspondence, whereas other receptors, and all transporters, show no expression-density correspondence. Since group-averaged measures of gene expression and receptor/transporter density come from disjoint samples of participants, we hypothesized that genes with greater variability between participants would show weaker correlations with group-averaged neurotransmitter receptor and transporter densities. To test this, we used each gene’s differential stability, a measure of gene expression variability across the six donors (see *Methods* for details [36]). Fig. 3a reveals that genes with greater differential stability, and therefore less inter-subject variance, are generally more correlated with PET-derived receptor density (*r* = 0.82, *p* = 4 × 10^-6^). When we repeated the analyses in the autoradiography dataset, we find a similar trend (*r* = 0.74, *p* = 0.0001; Fig. 3b).

**Figure 3.**
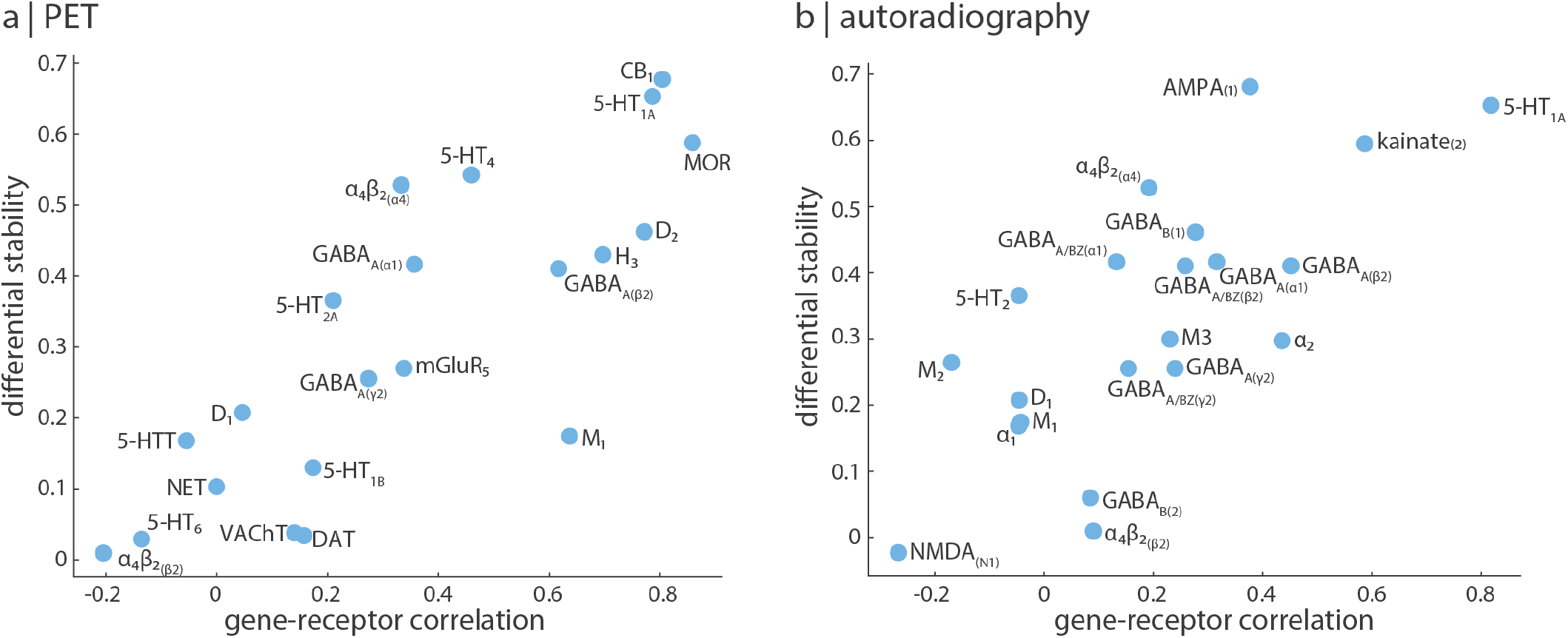
Differentially stable genes are more correlated with neurotransmitter receptor density. We find that differential stability, a measure of the variability of a gene’s expression across donors, is significantly correlated with gene-receptor correlations when using (a) PET-derived receptor/transporter densities (*r* = 0.82, *p* = 5 × 10^-6^) and (b) autoradiography-derived receptor densities (*r* = 0.74, *p* = 0.0001). Labels are neurotransmitter receptors/transporters, and suffixes in parentheses are relevant subunits.

The previous analyses were conducted across the full left hemisphere, but it is possible that gene-receptor/transporter coupling is variable across cortex. Since receptor distributions covary with cognitive functional activations along a sensory-fugal gradient [27, 34, 35], we tested whether gene expression is correlated with receptor densities across the four Mesulam classes of laminar differentiation (Fig. 4; [51, 61]). This analysis was done using the 111-region left hemisphere parcellation to ensure sufficient data observations in each laminar class. Interestingly, we find that receptors whose density is highly correlated with gene expression may show poor association in specific classes. Indeed, 5-HT_1A_ and D_2_ receptor density is highly correlated with expression in all laminar classes except idiotypic, which includes primary sensory-motor regions. The remaining receptors and transporters show large expression-density correlation variability across laminar classes. For example, 5-HT_6_ is consistently poorly correlated with *HTR6* gene expression no matter the laminar class (|*r*| < 0.13), whereas the GABAA receptor is highly correlated with *GABRB2* expression in idiotypic areas (*r* = 0.62) but weakly correlated in heteromodal and paralimbic areas (*r*_plmb_ = –0.10, *r*_het_ = 0.02). Meanwhile, dopamine receptor D_1_ is positively correlated with *DRD1* expression in paralimbic areas (*r* = 0.30), negatively correlated in idiotypic areas (*r* = –0.45), and not correlated in intermediate areas (|*r*| < 0.15). Altogether, we find that the correspondence between gene expression and receptor density is itself variable across the cortex and differs between laminar classes and cognitive systems [60, 90].

**Figure 4.**
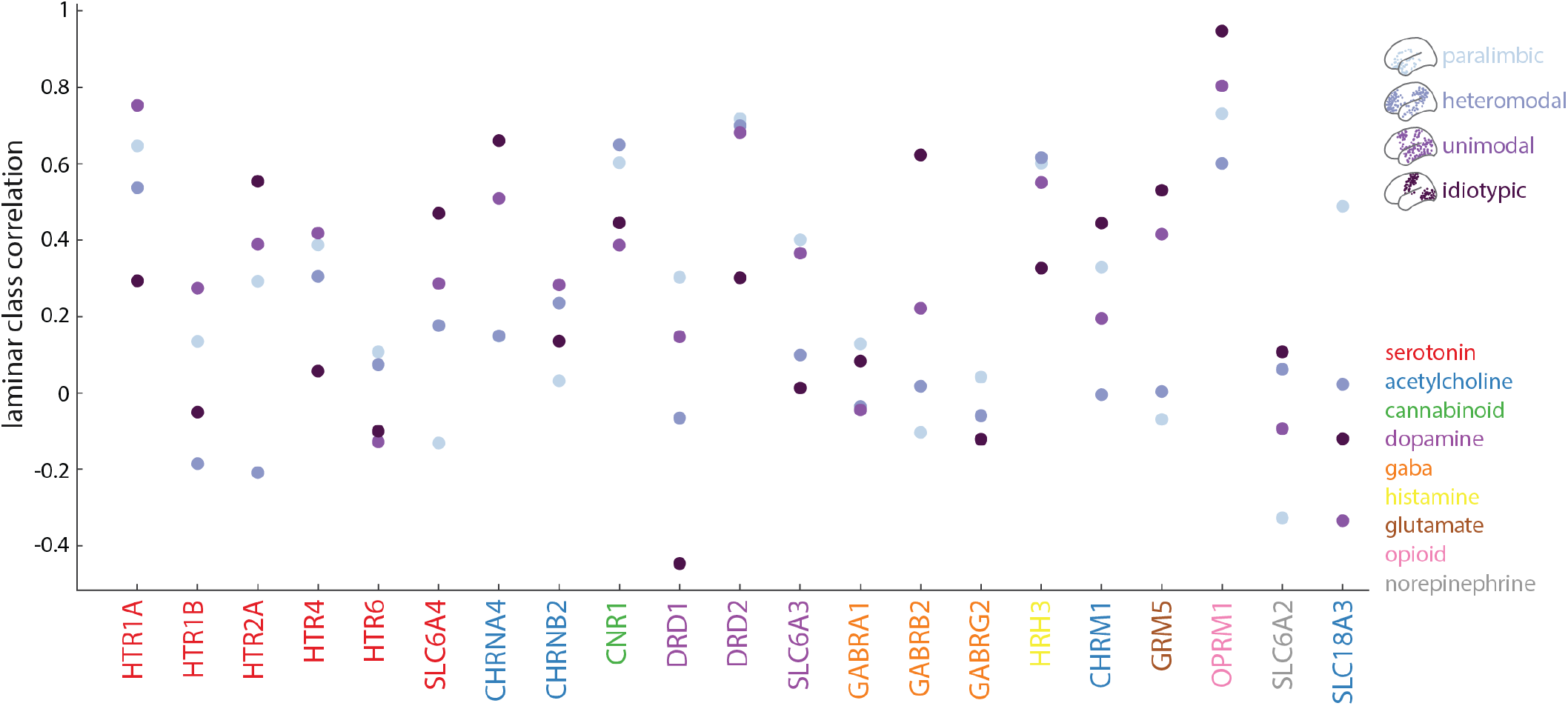
Gene-receptor/transporter correspondence varies across Mesulam classes of laminar differentiation. PET-derived receptor/transporter densities were correlated to gene expression within four classes of laminar differentiation [51, 61]. For each receptor and transporter, the density-expression correlation within each laminar class is plotted. Gene names are coloured by neurotransmitter system.

## DISCUSSION

Understanding how the chemoarchitecture of the brain modulates the link between structure and function requires accurate and comprehensive regional neurotransmitter receptor and transporter profiles. Here we formally test whether there is a correlation between gene expression and neurotransmitter receptor/transporter density, for a total of 27 unique neurotransmitter receptors, receptor binding sites, and transporters, from both PET images and autoradiographs. We find that only four receptors (5-HT_1A_, D_2_, CB_1_, and MOR) display a close spatial correspondence between gene expression levels and receptor densities. We therefore conclude that researchers should exercise caution when using gene expression as a proxy for receptor and transporter densities.

We note that the lack of correlation between protein levels and the levels of their coding mRNA is not unreasonable as there are many mechanisms that may affect the protein-mRNA correlation. First, levels of mRNA detected on the microarray do not take into account the transcript isoforms that can be produced from the same gene, nor the stability of the resulting mRNA, which are determined by mRNA modifications such as splicing [46, 86]. Second, the proportion of different cell types in a microarray sample may distort the gene expression-protein density correspondence due to differences in the proteome and transcriptome, including different splice variant expression [78, 89]. Third, studies in bacteria [45] and mice [40] have demonstrated that rate of protein synthesis also alters protein levels. Fourth, protein buffering dampens the effect of variations in gene expression levels, including an adaptation of protein turnover through protein degradation, and the modulated activity of protein transport machinery which determines the final subcellular localization of the protein [7, 46, 77, 87]. Altogether, the variation in the activity of these processes may contribute to the observed difference in levels of mRNA expression and protein abundance of receptors/transporters in the cortex.

Nonetheless, we do find a small subset of neurotransmitter receptors that demonstrate close gene transcription-receptor density relationships. One possible explanation is rooted in population variance of gene expression and receptor densities. Since both receptor densities and gene expression are averaged across participants, genes and receptors with low inter-subject variability across the population would be better captured by the group-averaged map used in the present analyses. This is supported by the fact that neurotransmitter receptor densities are more correlated with genes that are more stable across donors (Fig. 3). A second nonexclusive explanation is that the steps between gene transcription and membrane insertion are potentially more preserved for specific neurotransmitter receptors. Indeed, the correspondence between gene expression and receptor density would depend on localization of the mRNA due to differences in ribosome and tRNA availability [12], as well as protein turnover rates at the location the receptor is expressed [13]. Additionally, receptor systems that are phylogenetically older, involved in functions that require rigid stimulus-response relationships, or more fundamental to the organizational principles of the brain, may demonstrate more robust translation that manifest as close expression-density associations. For example, in the mouse cortex, gene expression for certain interneuron cell types are closely aligned with gradients of cortical organization [30]. Interestingly, the gene that most recapitulates this organizational feature (*Pvalb*) is highly correlated to its corresponding protein’s density (parvalbumin; Spearman *r* = 0.95), whereas *Sst*—an interneuron marker that is poorly aligned with cortical organization—is poorly correlated with somatostatin density (Spearman *r* = 0.24, *p* = 0.1) [28].

In the present report, we caution against the use of gene expression as a proxy for receptor densities, and recommend using techniques that more directly capture receptor distributions such as PET or autoradiography. However, we note that both PET and autoradiography have specific challenges. For example, PET tracers are generally only sensitive to receptors on the cell surface, but neurotransmitter receptors are also found within the cell [75]. Furthermore, PET does not directly measure density, is sensitive to in-scanner motion, and may demonstrate non-specific binding [26]. Meanwhile, autoradiographs more directly measure receptor densities but are only acquired post-mortem, are more expensive and labor intensive, and measure densities in discrete brain sections [92]. Although it is encouraging that we find consistent results across PET- and autoradiography-derived receptor densities, future research is needed to accurately and comprehensively measure neurotransmitter receptor densities throughout the brain.

Our results build on previous work that explores the expression-density relationship of specific neurotransmitter systems or receptors, such as the serotonergic system [10, 43], the GABA_A_ receptor [58], and the opioid system [43, 69], using PET and/or autoradiography-derived density. The present report comprehensively investigates the expression-density correspondence for 27 unique neurotransmitter receptors, receptor binding sites, and transporters across 9 different neurotransmitter systems using both PET and autoradiography measurements. Here we note some consistencies and inconsistencies across findings. Consistencies include: (1) high correlation between 5-HT_1A_ density and *HTR1A* expression [10, 43, 69], (2) weaker associations for other serotonergic receptors [10], (3) high correlation between MOR and *OPRM*_1_ [43], and (4) positive correlation between GABAA density and *β*_2_ subunit expression, but negative correlation between GABAA density and *γ*_1_ subunit expression—although we do not find that these relationships are significant after correcting for multiple comparisons [58]. On the other hand, we find no association between GABAA density and the expression of *α*_1_ and *γ*_2_ main channel subunits, unlike that reported in [58]. Additionally, [69] find that the PET tracer dipenorphine, which binds to all three opioid subtypes (*δ, κ*, and *μ*), is not correlated with the expression of any opioid subtype. Meanwhile, we find a strong correlation between MOR density and *OPRM1* expression, and presume that expression-density relationships are specific to single receptors and not generalizable across receptors in the same neurotransmitter system. We note that these inconsistencies are likely related to the gene normalization method, a key processing step with large effects on estimated gene expression [47].

We close with some methodological considerations when working with PET, autoradiography, and gene expression datasets. First, gene expression estimates are derived from only six post-mortem human brains. Although the Allen Human Brain Atlas is a state-of-the-art dataset of microarray gene expression, more comprehensive datasets are necessary to confirm gene expression levels. Second, measures of neurotransmitter receptor/transporter densities and gene expression are acquired in different individuals, so we are not able to make conclusions on the correspondence between gene expression and receptor/transporter density in the same cell tissue. Third, due to the relatively coarser resolution of PET, and the incomplete spatial coverage of autoradiography, main analyses were conducted in a parcellation of only 33–34 left hemisphere cortical brain regions. Replication in a finer parcellation for the PET receptor data do show similar results (Fig. S3), as well as high receptor density correlation between hemispheres, but high resolution whole-brain gene-receptor/transporter analyses should be conducted in future work.

In summary, we find that the expression of specific receptor/transporter-coding genes can generally not be used to estimate neurotransmitter receptor and transporter density. We only find a correspondence between gene expression and receptor density for 5-HT_1A_, D_2_, CB_1_, and MOR. Future efforts to map neurotransmitter receptor and transporter profiles to brain structure and function should verify the expression-density association when using microarray gene expression in place of receptor and transporter density.

## Acknowledgments

We thank Kelly Smart, Sylvia Cox, Yanjun Wu, Jean-Dominique Gallezot, Étienne Aumont, Stijn Servaes, Stephanie G. Scala, Jonathan M. DuBois, Gleb Bezgin, Taylor W. Schmitz, R. Nathan Spreng, Jean-Paul Soucy, Synthia Guimond, Jarmo Hietala, Marc-André Bédard, Marco Leyton, Eliane Kobayashi, Pedro Rosa-Neto, and Richard E. Carson for collecting and sharing the PET data presented here. We thank Vincent Bazinet, Zhen-Qi Liu, Filip Milisav, Laura Suarez, Bertha Vazquez-Rodriguez, and Mingze Li for their comments and suggestions on the manuscript. This research was undertaken thanks in part to funding from the Canada First Research Excellence Fund, awarded to McGill University for the Healthy Brains for Healthy Lives initiative. BM acknowledges support from the Natural Sciences and Engineering Research Council of Canada (NSERC Discovery Grant RG-PIN #017-04265) and from the Canada Research Chairs Program. JYH acknowledges support from the Helmholtz International BigBrain Analytics & Learning Laboratory, the Natural Sciences and Engineering Research Council of Canada, and the Fonds de reserches de Québec. The funders had no role in study design, data collection and analysis, decision to publish or preparation of the manuscript.

---

**Figure S1.**
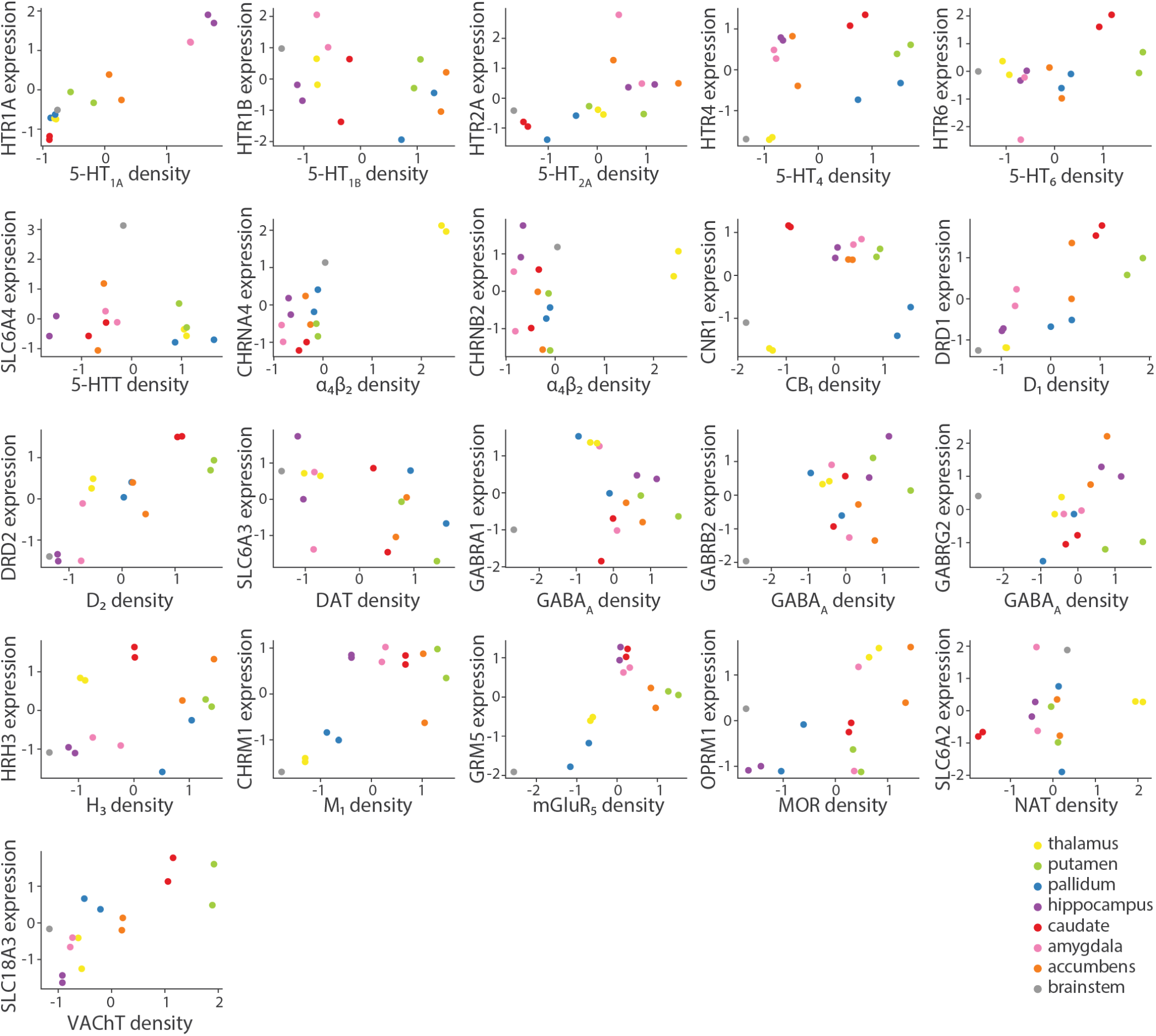
Correspondence between gene expression and receptor/transporter density in the subcortex. PET receptor/transporter densities and microarray gene expression was parcellated into 15 subcortical regions and correlated.

**Figure S2.**
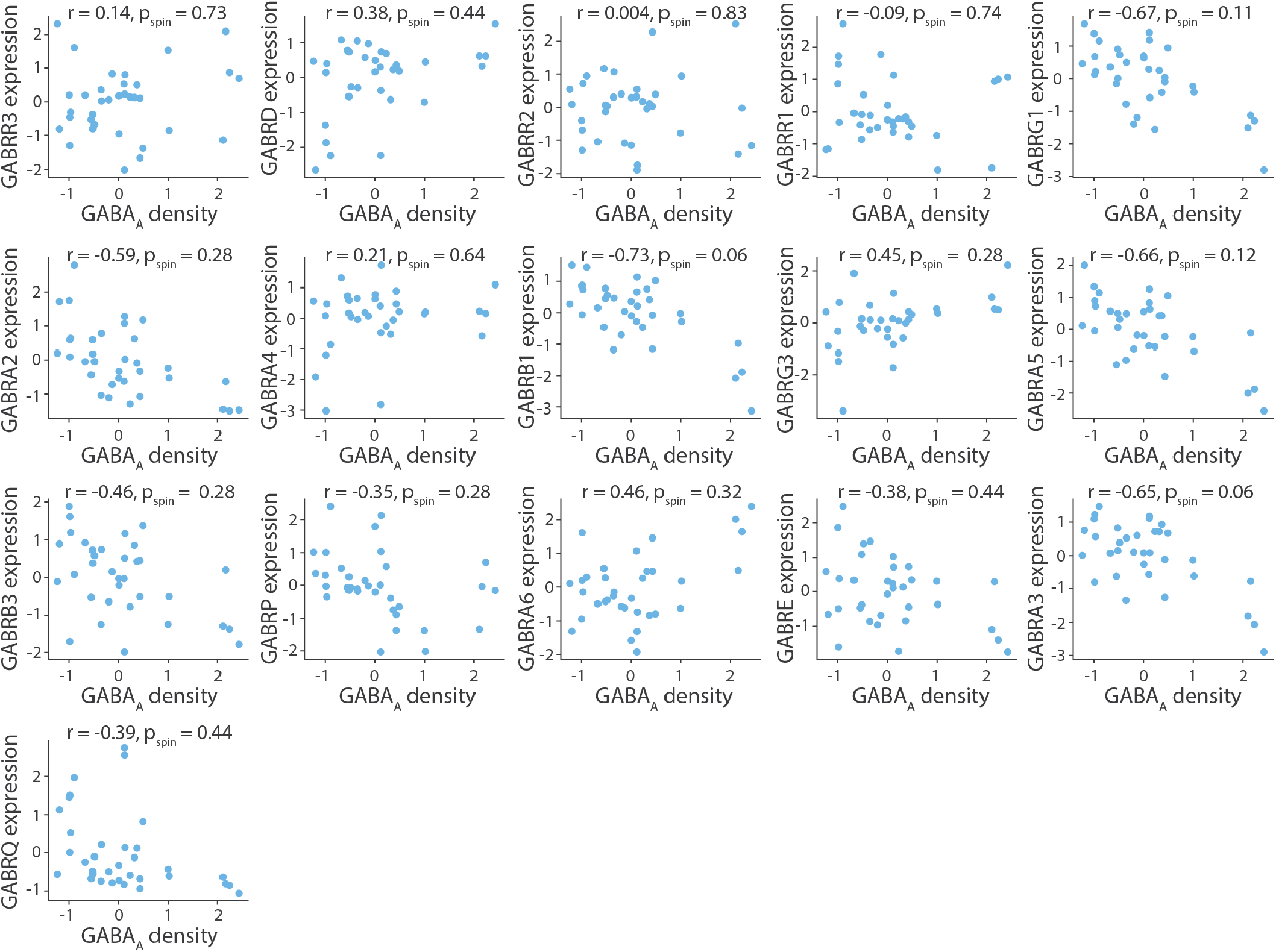
Density-expression association for the remaining sixteen GABAa subunits. Density-expression association for the remaining sixteen GABAa subunits that do not comprise the main channel show that the expression of four subunits are strongly negatively correlated (*r* < –0.6) with receptor density (*γ*_1_, *β*_1_, *α*_5_, *α*_3_), although the relationship is not significant after correcting for multiple comparisons [11]. All receptor densities are z-scored.

**Figure S3.**
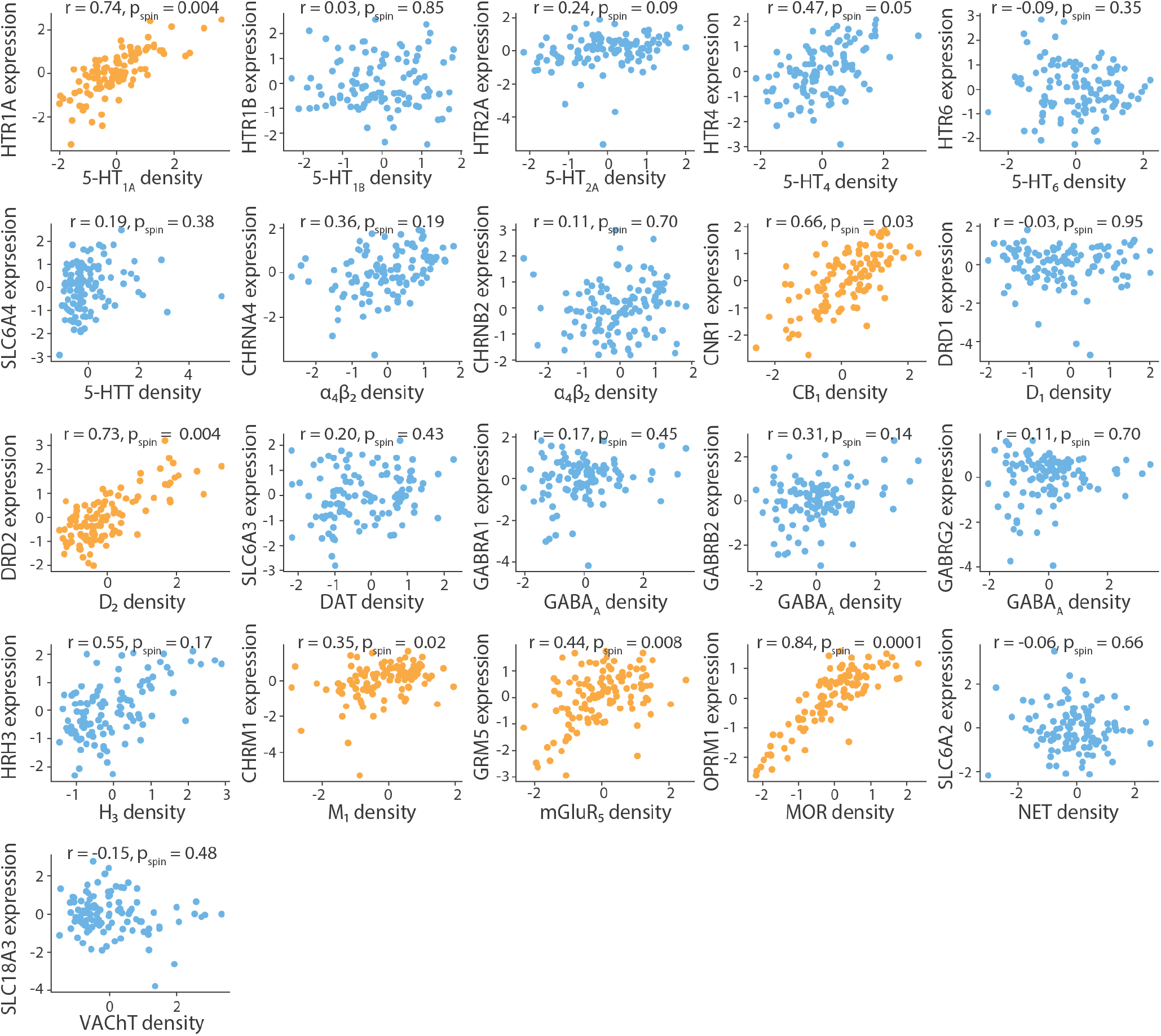
Replication in a 111-node parcellation. PET receptor/transporter densities and gene expression levels were parcellated into a 111-node cortical left hemisphere parcellation. Yellow scatter plots indicate significant density-expression correspondence, against an autocorrelation-preserving null model. All receptor and transporter densities are z-scored.

**TABLE S1.** Neurotransmitter receptors and transporters included in analyses. BP_ND_ = non-displaceable binding potential; V_T_ = tracer distribution volume; B_max_ = density (pmol/ml) converted from binding potential (5-HT) or distributional volume (GABA) using autoradiography-derived densities; SUVR = standard uptake value ratio. Neurotransmitter receptor maps without citations refer to unpublished data. Refer to [35] for more details. Asterisks indicate transporters.

| Receptor/<br>transporter | Neurotransmitter | Tracer | Measure | N | Age | References |
| --- | --- | --- | --- | --- | --- | --- |
| D<sub>1</sub> | dopamine | [<sup>11</sup>C]SCH23390 | BP<sub>ND</sub> | 13 | 33 ± 13 | Kaller et al., 2017 [41] |
| D<sub>2</sub> | dopamine | [<sup>11</sup>C]FLB-457 | BP<sub>ND</sub> | 37 | 48.4 ± 16.9 | Smith et al., 2019 [71, 83] |
| D<sub>2</sub> | dopamine | [<sup>11</sup>C]FLB-457 | BP<sub>ND</sub> | 55 | 32.5 ± 9.7 | Sandiego et al., 2015 [71, 72, 81, 83, 88] |
| DAT* | dopamine | [<sup>123</sup>I]-FP-CIT | SUVR | 174 | 61 ± 11 | Dukart et al., 2018 [24] |
| NET* | norepinephrine | [<sup>11</sup>C]MRB | BP<sub>ND</sub> | 77 | 33.4 ± 9.2 | Ding et al., 2010 [9, 18, 22, 70] |
| 5-HT<sub>1A</sub> | serotonin | [<sup>11</sup>C]WAY-100635 | BP<sub>ND</sub> | 36 | 26.3 ± 5.2 | Savli et al., 2012 [74] |
| 5-HT<sub>1B</sub> | serotonin | [<sup>11</sup>C]P943 | BP<sub>ND</sub> | 65 | 33.7 ± 9.7 | Gallezot et al., 2010 [5, 31, 50, 54, 55, 63, 73] |
| 5-HT<sub>1B</sub> | serotonin | [<sup>11</sup>C]P943 | BP<sub>ND</sub> | 23 | 28.7 ± 7.0 | Savli et al., 2012 [74] |
| 5-HT<sub>2A</sub> | serotonin | [<sup>11</sup>C]Cimbi-36 | B<sub>max</sub> | 29 | 22.6 ± 2.7 | Beliveau et al., 2017 [10] |
| 5-HT<sub>4</sub> | serotonin | [<sup>11</sup>C]SB207145 | B<sub>max</sub> | 59 | 25.9 ± 5.3 | Beliveau et al., 2017 [10] |
| 5-HT<sub>6</sub> | serotonin | [<sup>11</sup>C]GSK215083 | BP<sub>ND</sub> | 30 | 36.6 ± 9.0 | Radhakrishnan et al., 2018 [66, 67] |
| 5-HTT* | serotonin | [<sup>11</sup>C]DASB | B<sub>max</sub> | 100 | 25.1 ± 5.8 | Beliveau et al., 2017 [10] |
| α<sub>4</sub>β<sub>2</sub> | acetylcholine | [<sup>18</sup>F]flubatine | V<sub>T</sub> | 30 | 33.5 ± 10.7 | Hillmer et al., 2016 [4, 38] |
| M<sub>1</sub> | acetylcholine | [<sup>11</sup>C]LSN3172176 | BP<sub>ND</sub> | 24 | 40.5 ± 11.7 | Naganawa et al., 2021 [56] |
| VACHT* | acetylcholine | [<sup>18</sup>F]FEOBV | SUVR | 4 | 37 ± 10.2 | PI: Lauri Tuominen & Synthia Guimond |
| VACHT* | acetylcholine | [<sup>18</sup>F]FEOBV | SUVR | 18 | 66.8 ± 6.8 | Aghourian et al., 2017 [1] |
| VACHT* | acetylcholine | [<sup>18</sup>F]FEOBV | SUVR | 5 | 68.3 ± 3.1 | Bedard et al., 2019 [8] |
| VACHT* | acetylcholine | [<sup>18</sup>F]FEOBV | SUVR | 3 | 66.6 ± 0.94 | PI: Taylor W. Schmitz & R. Nathan Spreng |
| mGluR<sub>5</sub> | glutamate | [<sup>11</sup>C]ABP688 | BP<sub>ND</sub> | 73 | 19.9 ± 3.04 | Smart et al., 2019 [82] |
| mGluR<sub>5</sub> | glutamate | [<sup>11</sup>C]ABP688 | BP<sub>ND</sub> | 22 | 67.9 ± 9.6 | PI: Pedro Rosa-Neto |
| mGluR<sub>5</sub> | glutamate | [<sup>11</sup>C]ABP688 | BP<sub>ND</sub> | 28 | 33.1 ± 11.2 | DuBois et al., 2016 [23] |
| GABA<sub>A/BZ</sub> | GABA | [<sup>11</sup>C]flumazenil | B<sub>max</sub> | 16 | 26.6 ± 8 | Nørgaard et al., 2021 [58] |
| H<sub>3</sub> | histamine | [<sup>11</sup>C]GSK189254 | V<sub>T</sub> | 8 | 31.7 ± 9.0 | Gallezot et al., 2017 [32] |
| CB<sub>1</sub> | cannabinoid | [<sup>11</sup>C]OMAR | V<sub>T</sub> | 77 | 30.0 ± 8.9 | Normandin et al., 2015 [25, 57, 59, 68] |
| MOR | opioid | [<sup>11</sup>C]carfentanil | BP<sub>ND</sub> | 204 | 32.3 ± 10.8 | Kantonen et al., 2020 [42] |

**TABLE S2.**
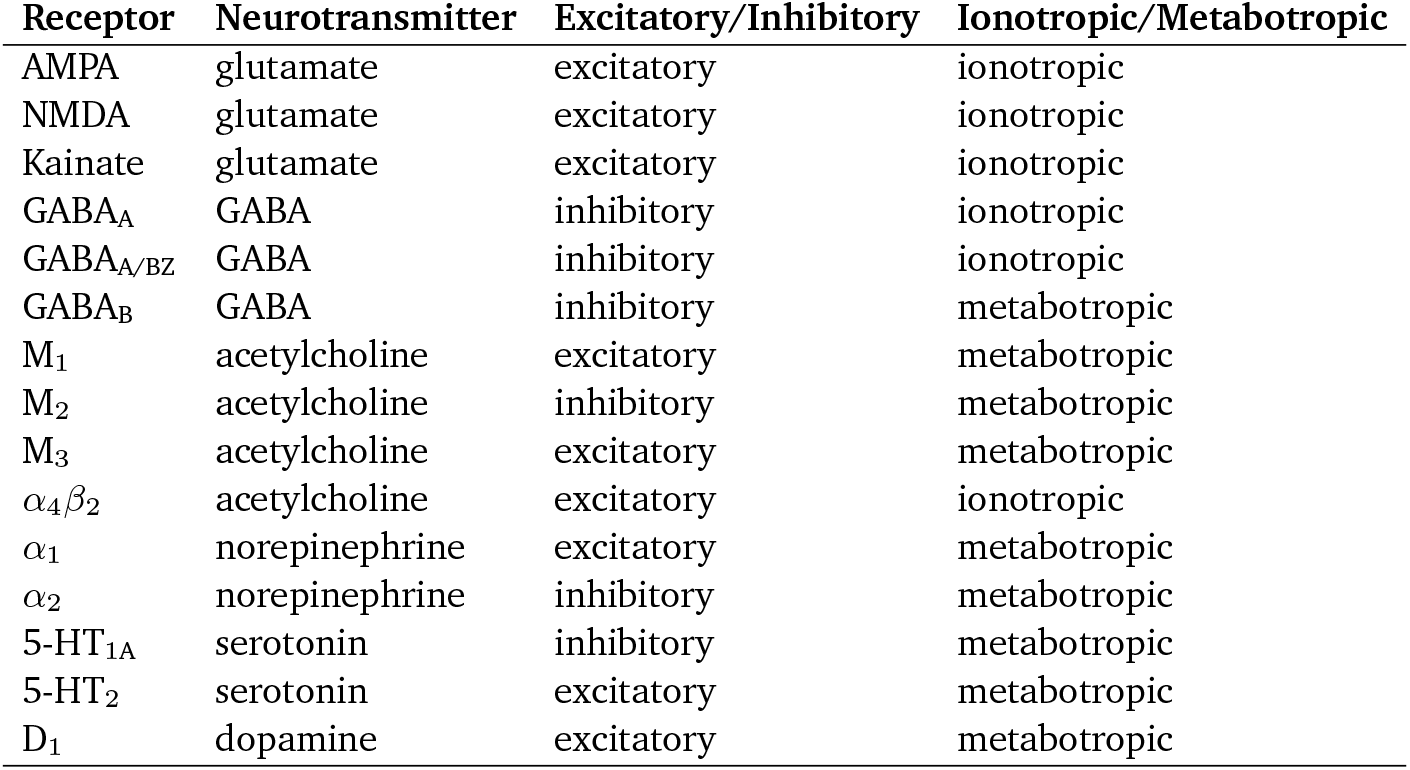
Neurotransmitter receptors included in the autoradiography dataset.

